# Intrinsic furin-mediated cleavability of the spike S1/S2 site from SARS-CoV-2 variant B.1.1.529 (Omicron)

**DOI:** 10.1101/2022.04.20.488969

**Authors:** Bailey Lubinski, Javier A. Jaimes, Gary R. Whittaker

## Abstract

The ability of SARS-CoV-2 to be primed for viral entry by the host cell protease furin has become one of the most investigated of the numerous transmission and pathogenicity features of the virus. SARS-CoV-2 The variant B.1.1.529 (Omicron) emerged in late 2020 and has continued to evolve and is now present in several distinct sub-variants. Here, we analyzed the “furin cleavage site” of the spike protein of SARS-CoV-2 B.1.1.529 (Omicron variant) *in vitro*, to assess the role of two key mutations (spike, N679K and P681H) that are common across all subvariants compared to the ancestral B.1 virus and other notable lineages. We observed significantly increased intrinsic cleavability with furin compared to an original B lineage virus (Wuhan-Hu1), as well as to two variants, B.1.1.7 (Alpha) and B.1.617 (Delta) that subsequently had wide circulation. Increased furin-mediated cleavage was attributed to the N679K mutation, which lies outside the conventional furin binding pocket. Our findings suggest that B.1.1.529 (Omicron variant) has gained genetic features linked to intrinsic furin cleavability, in line with its evolution within the population as the COVID-19 pandemic has proceeded.

## Introduction

The so-called furin cleavage site has become one of the most established transmission and pathogenicity factors of the SARS-CoV-2 spike protein (1), recently reviewed in (2). A structural loop containing a _682-_RRAR-_685_ amino acid motif was found to be present in the original SARS-CoV-2 isolates (3), with this “furin cleavage site” (FCS) located between the spike receptor-binding (S1) and fusion (S2) domains (S1/S2). An “FCS” has been subsequently found in all circulating variants but has evolved during the course of the pandemic, with the upstream proline (P_681_) residue being replaced by histidine (H) and arginine (R) in several variants of concern—notably B.1.1.7 (Alpha) and B.1.167 (Delta), respectively. This loop/FCS is missing from all other SARS-like viruses (Sarbecoviruses, betacoronavirus lineage B), and is actively de-selected upon adaptation of SARS-CoV-2 to common cell culture systems (e.g., Vero cells); see ref (4) as an example.

In late November 2021, a distinct SARS-CoV-2 variant emerged (B.1.1.529) that had an unusual cluster of mutations, especially in the spike gene (5, 6). The original variant was named “Omicron” by the World Health Organization, with the original lineage subsequently termed BA.1. There was an unprecedented and rapidly disseminated series of studies on this variant, with early studies describing quite different infection properties compared to all other variants that had emerged by that time. In summary, BA.1 was generally shown to be less fusogenic, and with lowered S1/S2 cleavage and TMPRSS-2 dependence than the other variants (7–14)—especially in comparison to B.1.617.2 (Delta) the predominant variant at the time of Omicron emergence. Pathogenicity in animal models was also reduced with Omicron BA.1. Many details of the cleavage-activation process of Omicron remain to be resolved, especially when considering that multiple sub-variants including BA.2, BA.4/5, BA.2.12.1 and BA.2.75 have now emerged—with BA.5 in particular seeming to show infection properties more in line with Delta and the original B.1 strain, including increased syncytia formation, TMPRSS-2 usage and pathogenesis (15, 16). Despite the multiple changes in the spike gene between the various Omicron sub-variants, the S1/S2 “furin” cleavage site has remained unchanged.

The furin cleavage site for proteins was initially described as a four amino acid pattern _P4_-R-X-[K/R]-R-_P1_ (17), with the cleavage occurring between the P1 and P1’ residues, as defined for protease substrates (18). This motif had become established as a prototype FCS, although the minimal residues for furin cleavage are generally defined as _P4_-R-X-X-R-_P1_ (19). Notably, the sequence context of the four amino acid “FCS” readily modulates cleavability, and subsequent analysis has characterized the furin cleavage motif as a 20 amino acid sequence running from position P14 to P6’—having a “core” region (P6-P2’), along with two flanking solvent accessible regions, P14-P7 and P3’-P6’ (20, 21).

The emergence of Omicron was reminiscent of that of the Alpha variant approximately one year earlier; both were notable for a suddenly increased accumulation of mutations, and both have been linked to long-term infection of individuals with persistent SARS-CoV-2 infection (22). Both Alpha and Omicron contain the same P681H mutation in the FCS (P5 residue), but Omicron contains an additional mutation N679K, which is the P7 residue for furin-mediated cleavage. While within the 20 amino acid sequence comprising a furin cleavage motif, P7 is outside of the core region comprising the established furin binding pocket (20). N679K introduces an additional basic residue into the FCS, which may result in a more “polybasic” sequence and increase furin-mediated cleavage.

Here, we analyze the intrinsic furin-mediated cleavage of the B.1.1.529 (Omicron) spike “FCS” in comparison to other SARS-CoV-2 variants, and discuss the impact of our observed increased cleavage on the COVID-19 pandemic and virus evolution.

## Methods

Fluorogenic peptide cleavage assays were performed as described previously (23), with minor modifications as described below. Reactions were carried out in a 96 well plate with a 100 μl volume consisting of cleavage buffer, protease, and fluorogenic peptide. Peptides were labeled with an MCA/DNP fluorescence resonance energy transfer (FRET) pair. Peptide sequences are listed in Table 1. Trypsin catalyzed reactions acted as a positive control, where 0.8 nM/well TPCK trypsin was diluted in PBS buffer. For furin catalyzed reactions, 1 U/well recombinant furin was diluted in a buffer consisting of 20 mM HEPES, 0.2 mM CaCl2, and 0.2 mM β-mercaptoethanol, at pH 7.5. A negative control consisted of peptide suspended in buffer for furin cleavage with no protease added (not shown). Fluorescence emission was measured once per minute for 60 continuous minutes using a SpectraMax fluorometer (Molecular Devices) at 30°C with an excitation wavelength of 330 nm and an emission wavelength of 390 nm. V_max_ was calculated by fitting the linear rise in fluorescence to the equation of the line.

**Table 1.**

| Peptide Name | Peptide Sequence |
| --- | --- |
| Wuhan Hu 1 | MCA-TQTNSPRRARSVA-Lys(DNP) |
| Alpha | MCA-TQTNSHRRARSVA-Lys(DNP) |
| Delta | MCA-TQTNSRRRARSVA-Lys(DNP) |
| Omicron | MCA-TQTKSHRRARSVA-Lys(DNP) |
| N679K P681R | MCA-TQTKSRRRARSVA-Lys(DNP) |
| N679K P681 | MCA-TQTKSPRRARSVA-Lys(DNP) |

## Results

As an initial bioinformatic approach to assess biochemical function, we utilized the PiTou (20) and ProP (24) protein cleavage prediction tools, comparing the spike proteins from B.1.1.529 to B.1.1.7, B.1.617 and the prototype SARS-CoV-2 sequence from the A.1 and B.1 lineages (e.g., Wuhan-Hu-1), as well as to MERS-CoV and selected other human respiratory betacoronaviruses that have identifiable furin cleavage sites (HCoV-HKU1 and HCoV-OC43). PiTou is expected to be a more accurate prediction tool as it utilizes biological knowledge-based cumulative probability score functions, combined with a hidden Markov model, and is reported to have a sensitivity of 96.9% and a specificity of 97.3% (20). ProP is a widely-used, but simpler, algorithm based on a pure machine learning neural network, with a reported sensitivity of 94.7% and a specificity of only 83.7 % (20)

For SARS-CoV-2 spike, only the ProP algorithm predicted an increase in the furin cleavage for the B.1.1.529 lineage, compared to other lineages (Figure 1). Pitou calculated B.1.1.529 as having the same score as B.1.1.7. As expected, both algorithms predicted MERS-CoV to have a relatively low furin cleavage score, with HCoV-HKU1 and HCoV-OC43 showing relatively high furin cleavage scores using both approaches.

**Figure 1:**
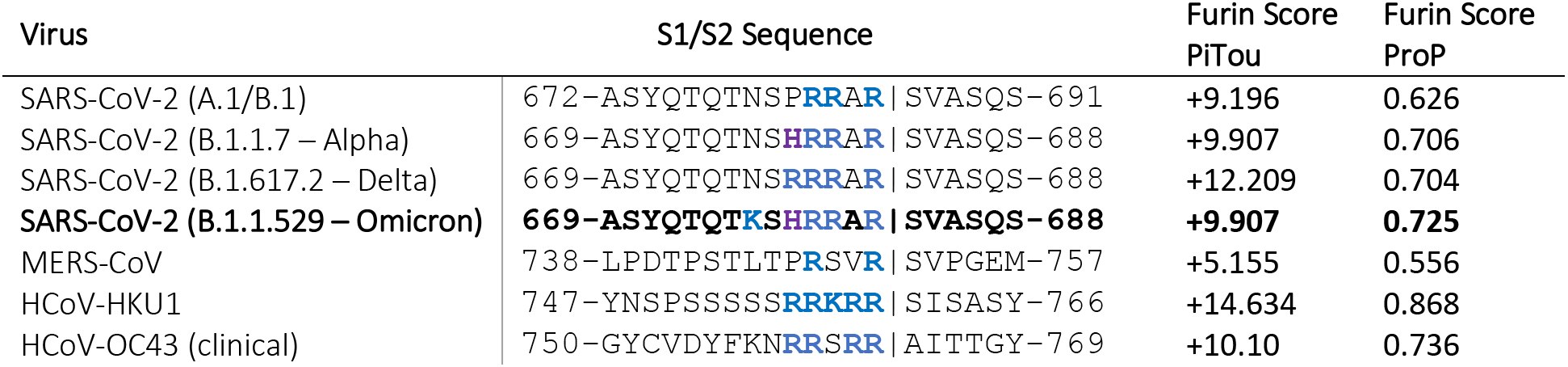
Predicted furin cleavage scores of the S1/S2 region of selected coronavirus spike proteins.

**Figure 2:**
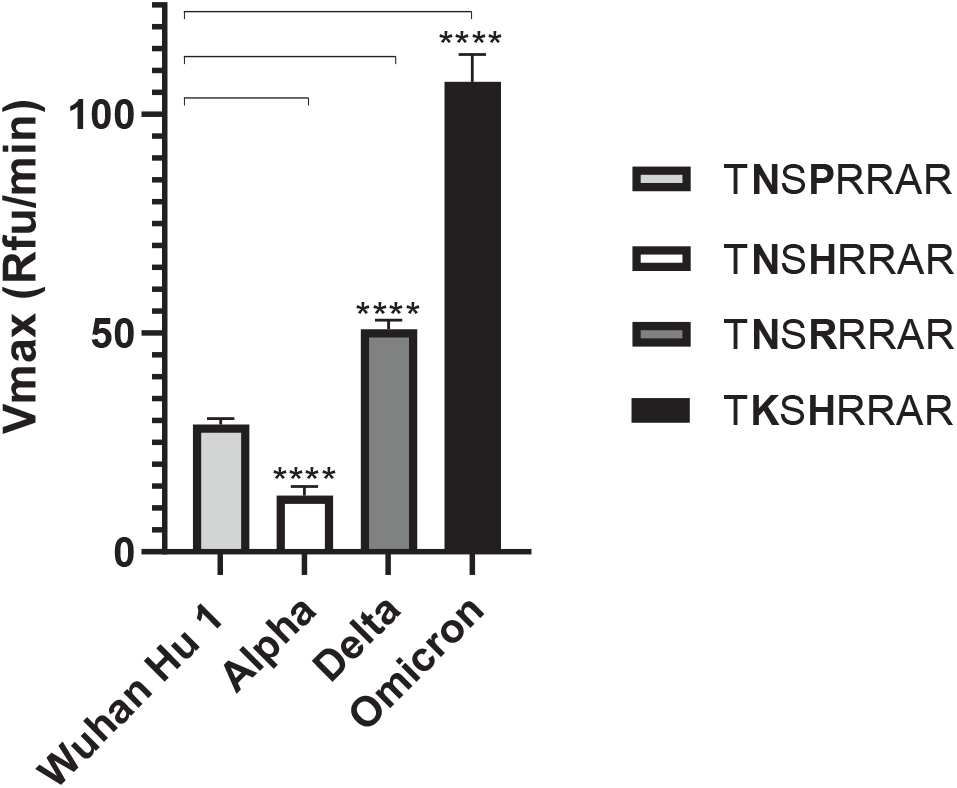
Fluorogenic cleavage assays of synthetic peptides labeled with an MCA/DNP FRET pair. Peptides were designed and synthesized to reproduce the S1/S2 sites of the spike protein of selected SARS-CoV-2 variants. Peptides were incubated with fUrin, fluorescence was monitored for 1 h and the V_max_ calculated. Error bars denote the standard error of the mean (SEM) (n=9). Two-tailed unpaired T tests were performed comparing Wuhan Hu 1 to each variant, with significance marked by asterisks. (****, p < 0.0001)

**Figure 3:**
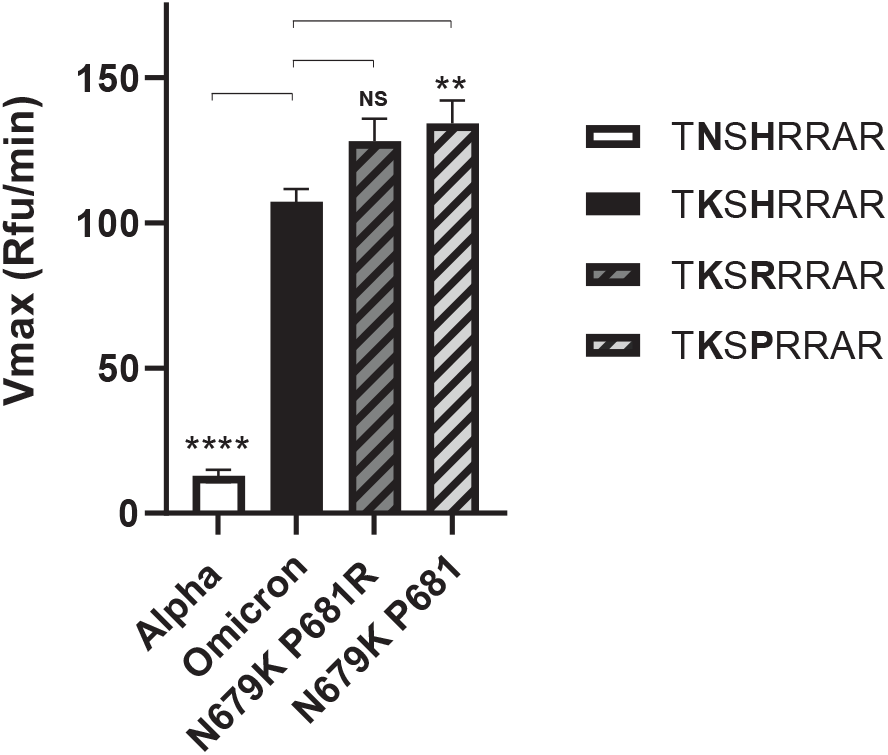
Fluorogenic cleavage assays of synthetic peptides labeled with an MCA/DNP FRET pair. Peptides were designed and synthesized to reproduce the S1/S2 site of the SARS-CoV-2 spike protein, examining the effect of introducing a lysine (K) residue at P7 (position 679) along with different established amino acids present at P5 (position 681). Peptides were incubated with furin, fluorescence was monitored for 1 h and the V_max_ calculated. Error bars denote the standard error of the mean (SEM) (n=9). Two-tailed unpaired T tests were performed comparing Omicron to each peptide, with significance marked by asterisks. (NS, p = 0.0308; **, p = 0.0085; ****, p < 0.0001)

To directly examine the activity of furin on the SARS-CoV-2 A.23.1 S1/S2 site, we used a biochemical peptide cleavage assay to directly measure furin cleavage activity *in vitro* (25). The specific peptide sequences used here are described in Table 1.

We found a noticeable increase in cleavability for Omicron compared to B.1 and all other variants tested, including Delta. With the peptides tested cleavage of Alpha (P681H) was actually decreased compared to B.1, indicating that the P7 K residue has a major impact on furin-mediated cleavage, despite it being outside the proposed “core” furin binding pocket. Trypsin was used as a control (see Appendix).

To explore this increased cleavage in more detail, we tested peptides with a P7 lysine residue (K) on the background of either B.1 and Delta FCS sequences (i.e., containing P681 and P681R). In both cases, the P7 K had a notable impact on cleavability, as with Omicron. The peptide with a P7 lysine residue on a B.1 background (P681) had a significantly higher cleavage rate when compared with Omicron, but how this increase in peptide cleavability translates to cleavability in the context of a full spike remains to be seen.

## Discussion

On its initial emergence SARS-CoV-2 B.1.1.529 (Omicron) showed surprising infection properties compared to other variants, including distinct cell tropism and cell-cell fusion, which were generally attributed to lowered S1/S2 “furin” cleavage and TMPRSS2-dependence (see Introduction). Conversely, it was also reported that the furin motif in the recombinant Omicron BA.1 spike ectodomain is more efficient than that in the prototypic spike ectodomain (26). These properties have since shifted with the emergence of subsequent sub variants (including BA.2, BA.4/5, BA.2.12.1 and BA.2.75).

In line with the most recent cell-cell fusion data on BA.5, our results show that SARS-CoV-2 B.1.1.529 (Omicron) has intrinsically high ability to be cleaved by furin at its spike S1/S2 position, despite early studies on BA.1 indicating the contrary—and despite the lowered score predicted by the Pitou FCS cleavage algorithm. In our study, *in vitro* peptide cleavage data are in line with the ProP FCS prediction tool, and with the data on recombinant S from Ye *et al*. (ref 26). Our data further suggest that N679K is a key mutation impacting SARS-CoV-2 spike protein function, especially when combined with additional changes at position 681 (R/H). While our data suggest that these changes are linked to furin processivity, other investigators have recently attributed altered functionality to other proteases, notably cathepsin G (27).

In addition to N679K/P681H is also notable B.1.1.529 (Omicron) has addition spike mutations such as H655Y and S375F that have been shown to allow altered spike cleavage-activation (28–30), with sub-variant BA.1 (but not BA.2) having ΔH69/V70 deletion in the N-terminal domain. All of these mutations, along with D614G (31), are likely to have an impact on cleavage efficiency and fusogencity for the intact spike protein. Our view is that SARS-CoV-2 B.1.1.529 (Omicron) has increased intrinsic S1/S2 spike cleavability, which is balanced by epistatic effects of other mutations to drive viral infection and cell tropism in a highly context-dependent manner.

It is noteworthy that mutations upstream of the S1/S2 cleavage motif, including Q675H/R, Q677 H/P as well as N679K have also occurred independently in many SARS-CoV-2 variants (32) and may confer more of a direct impact on furin processing, as they occur in the P9, P11 and P13 positions within the 20 amino acid furin cleavage motif. Despite overall mixed conclusions on Omicron spike cleavage, fusion and infection, our biochemical data support the hypothesis that SARS-CoV-2 is evolving toward increased inherent furin cleavability, with residue 679 (N-K) being a major site of selection.

## Acknowledgements

This work was funded in part by the National Institute of Health research grant R01AI35270 (to GW). We thank the global SARS-CoV-2 sequencing groups for their open and rapid sharing of sequence data and GISAID for providing an effective platform to make these data available, and all members of the Whittaker lab for helpful discussions.

## Appendix

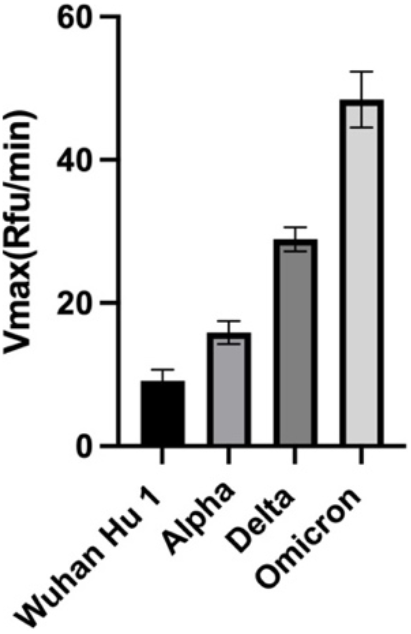

As a control, we also tested the ability of trypsin to cleave selected S1/S2 peptides. All variants were cleaved more effectively than for the prototype B.1 virus (Wuhan Hu 1), with B.1.1.529 (Omicron) showing the most cleavability.

